# The Aquaporin-4 expression and localization in the olfactory epithelium modulate the odorant-evoked responses and olfactory driven behavior

**DOI:** 10.1101/2024.09.03.610981

**Authors:** Donatella Lobraico, Pasqua Abbrescia, Maria Grazia Fioriello, Barbara Barile, Claudia Palazzo, Onofrio Valente, Grazia Paola Nicchia, Michele Dibattista, Antonio Frigeri

## Abstract

Aquaporin-4 (AQP4) is a water-selective channel expressed in glial cells throughout the central nervous system. It serves as the main water channel in the neuropil, and is involved in various physiological functions, ranging from regulating water homeostasis by adjusting cell volume to modulating neuronal activity. Different isoforms of AQP4 are expressed in glial-like cells known as sustentacular cells (SUSs) of the olfactory epithelium (OE). Interestingly, mice lacking all AQP4 isoforms exhibit impaired olfactory abilities. Hence, we aim to uncover the physiological role of two AQP4 isoforms, the perivascular AQP4ex isoform and the Orthogonal Array of Particle (OAP)-forming isoform (AQP4M23) in the OE. Primarily, we investigated the impact of AQP4 isoforms on OE protein expression, finding reduced levels of mature olfactory sensory neurons (OSNs) in mice lacking AQP4ex (AQP4ex-KO) or OAPs (OAP-null). Moreover, the reduced number of OSNs, SUSs, and globose basal cells (GBCs) suggests that AQP4 isoforms are involved in maintaining an optimal microenvironment in the OE, preserving the overall cell density. Then, we explored the role of AQP4 in modulating odorant-evoked responses through electro-olfactogram recordings, finding reduced odorant responses in mice lacking AQP4 isoforms. Olfactory ability assessments revealed deficits in odor-guided food-seeking test in AQP4ex-KO and OAP-null mice. Furthermore, AQP4ex-KO mice showed a reduced ability to discriminate between different odorants, while OAP-null mice were unable to recognize them as distinct. Overall, our data highlight the role of AQP4 isoforms in modulating neuronal homeostasis, affecting odorant-evoked responses and cell density in the OE. These results shed light on SUSs involvement in mediating these processes and establish a foundation for further understanding their role in controlling OE physiology.

## Introduction

Olfaction is one of the most ancient sensory systems, and its functional unit, the olfactory sensory neuron (OSN) is located in the olfactory epithelium (OE) (Buck et Axel, 1991; Niimura, 2012; Arguello et al., 2021). Since what was the first description of the OE, cell types other than OSNs were identified, such as the supporting cells (SUSs) (Allison, 1953). SUSs enwrap mature OSNs, suggesting they could provide mechanical, trophic and metabolic support and modulate the OSNs’ electrical activity (Morrison and Costanzo, 1992; Nomura et al., 2004; Liang, 2018; Hernandez-Clavijo et al., 2021). Indeed, SUSs modulate OSN activity by producing a *plethora* of neuromodulator molecules, such as NPY, an anorexigenic peptide, BDNF (brain-derived neurotrophic factor), and ATP, among others, that can modulate the odorant-evoked electrical response in OSNs by increasing the Ca^2+^ influx (Hansel, et al., 2001; Hayoz et al., 2012; Frontera, et al., 2015; Henriques, et al., 2019). Various modulatory activities promoted by SUSs are related to nucleotide signaling in the OE. SUSs express metabotropic P2Y purinergic receptors, and ATP induces intracellular Ca^2+^ influx in both SUSs and OSNs, modulating the odorant-evoked electrical response in OSNs (Hegg et al., 2003; Hegg et al., 2009; Dooley et al., 2011). SUSs can sustain OSN activity by releasing glucose into the mucus that covers the OE. Then, glucose is taken up into OSN cilia and metabolized to generate ATP to sustain olfactory transduction (Villar et al., 2017; Acevedo et al., 2019). Similarly to the astrocytes, the modulation activity could also be mediated by the gap junctions by offering cytoplasmic continuity and electrical coupling between OSNs and SUSs and providing a pathway for intercellular communication via Ca^2+^ and other signaling molecules (Rash et al., 2005; Vogalis et al., 2005a; Vogalis et al., 2005b; Hegg, et al., 2009). Therefore, the involvement of the SUSs in the aforementioned neuro-excitatory and/or neuro-modulatory phenomena makes them similar to the astrocytes or glial cells in peripheral organs like Müller cells in the retina. These cells adapt their dynamics, supporting and synchronizing local electrical activity (Newman, 1987; Gotow et Hashimoto, 1989; Parpura et al., 1994; Duan et al., 2003; Nedergaard et al., 2003; Jourdain et al., 2007; Rouach et al., 2008). Such plasticity is mediated by ion and water flux, relying on the water channel aquaporin-4 (AQP4) (Saadoun et al., 2005; Skucas et al., 2011; Li et al., 2012; Ciappelloni et al., 2019).

AQP4 expression has been reported in the mouse OE, including the basal cells (BCs), Bowman’s glands and SUSs (Nielsen et al., 1997b; Ablimit et al., 2006; Solbu and Holen, 2012).

AQP4 is the water channel mainly expressed in end-foot astrocyte processes and ependymal cells surrounding the ventricles (Frigeri et al., 1995; Nielsen et al., 1997a; Rash et al., 1998). It is involved in promoting water movements in the central nervous system (CNS), with a prominent role in determining blood-brain barrier (BBB) water permeability (Nicchia et al., 2004; Oshio et al., 2004; Papadopoulos et al., 2004). The water channel presents two major isoforms: a longer isoform starting with Met-1, named M1 (32 kDa), and a shorter isoform with a translation starting point at Met-23, named M23 (30 kDa), produced by alternative splicing called leaky scanning (Rossi et al., 2010). AQP4 may even undergo an alternative translational modification, called translational readthrough, providing an extension of roughly 29 amino acids at the C-terminus, thus generating the extended isoforms (AQP4ex) of either AQP4M23 (M23ex) and AQP4M1 (M1ex) (Loughran et al., 2014; De Bellis et al., 2017). Unlike what occurs in other AQPs, which assemble in homotetramers and then form intramembrane particles, the M1 and M23 isoforms are assembled in cell membranes as AQP4 heterotetramers (Lu et al., 1996; Furman et al., 2003). Freeze-fracture electron microscopy has shown square arrays of intramembrane particles in cell membranes expressing AQP4, known as Orthogonal Array of Particles (OAPs) (Landis and Reese, 1974; Crane, et al., 2009; Hirt et al., 2011; Jin et al., 2011). The ratio of M1 and M23 expression determines the size of the OAPs (Silberstein et al., 2004; Nicchia et al., 2010; Rossi et al., 2010). Moreover, AQP4ex is required to anchor OAPs at the perivascular astrocytic end-feet, and its absence induces rearrangement and redistribution of OAPs (Palazzo et al., 2019; Palazzo et al., 2020; Pati et al., 2022).

AQP4 is expressed in glial-like SUSs, even though little is known about its role in the OE, and nothing is known about the role of the AQP4ex isoforms in peripheral sensory organs, such as the OE. On the other hand, the involvement of the astrocytes and Müller cells in maintaining an optimum microenvironment in the extracellular space (ECS) through AQP4 is well known. Therefore, can SUSs be involved in similar roles of the glial cells in the OE? What is the contribution of AQP4’s isoforms? By taking advantage of AQP4ex-KO mice, specifically lacking AQP4ex-M1 and AQP4ex-M23 isoforms, and OAP-null mouse model, lacking AQP4M23 and AQP4ex-M23 isoforms, we explored the role of the OAPs in the OE and reported the localization of the newly described AQP4ex in the OE, and investigated its role in olfactory physiology and behavior.

## Results

### Localization of AQP4 and AQP4ex in the olfactory epithelium

The main water channel of the nervous system, AQP4, has different isoforms that have been described (Hasegawa et al., 1994; Jung, et al., 1994; De Bellis et al., 2017). The extended isoforms produced via translational readthrough have been shown to be crucial for the proper cellular compartmentalization of the water channel in glial cells of different districts in the brain (De Bellis et al., 2017; Palazzo et al., 2019; Palazzo et al., 2020). It is unknown whether the extended isoforms are expressed in the sensory organs and their functional roles.

In the OE, by using an antibody that detects all the AQP4 isoforms we found that all of them are localized on the basolateral membrane of SUSs, and appear to wrap OSN membranes in the olfactory epithelium, where the OSN cell bodies are located (Fig. 1 a-c). AQP4 expression spread across the OE, mutually exclusive with olfactory marker protein (OMP) staining, a marker for mature OSNs, demonstrating that AQP4 is not expressed in the OSNs (Fig. 1 a-c). Furthermore, AQP4 staining is intense at the bottom of the OE above the basal lamina, where the basal stem cells reside (Fig. 1 a). We found that AQP4 is also expressed in the cells below the basal lamina, most likely olfactory ensheathing glial cells (OECs) in close contact with OSN axon bundles (Fig. 1 a-c). By using an antibody specifically detecting the extended isoforms of AQP4 (AQP4ex, see methods) we observed that AQP4ex has higher expression in the basal part of the OE, above and below the basal membrane. Indeed, AQP4ex seems expressed in the basal cells, even though we cannot completely rule out that the signal could resemble the compartmentalization of AQP4ex in the basal foot of the supporting cells (Fig. 1 d-f). The nuclei of SUSs and basal cells were identified with an antibody against the transcription factor Sox2 (Guo, et al., 2010). The SUS nuclei are lined up in the apical part of the OE, and AQP4 staining is surrounding the cells’ nuclei, indicating the presence of AQP4 channels in the lateral membranes of SUS cells. SOX2^+^ basal cells are surrounded by AQP4 (Fig. 1 g-i), thus strongly suggesting that basal cells express AQP4 on their cell membrane.

**Figure 1.**
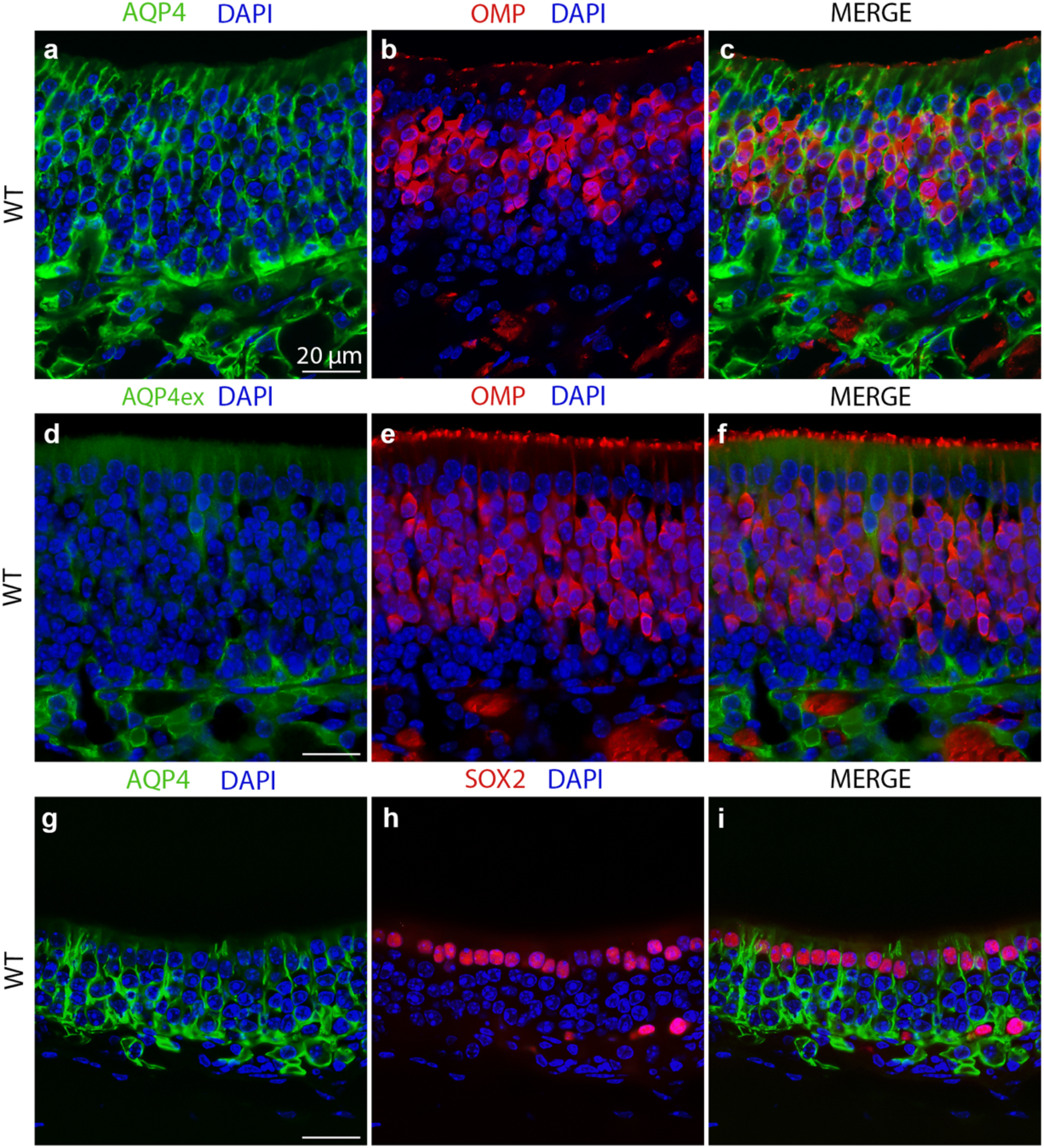
AQP4 is expressed on the basolateral membrane of the sustentacular cells. (a, b, c) High-magnification confocal micrographs of coronal sections from wild-type mouse olfactory epithelium showing AQP4 expression pattern. OMP staining does not colocalize with AQP4. AQP4 is also localized beneath the basal lamina. (d, e, f) AQP4ex shows the same localization as AQP4. (g, h, i) AQP4 staining in the basolateral membrane of the sustentacular cells stained with Sox2. Sox2^+^ cell nuclei are surrounded by AQP4.

These findings are further confirmed by the AQP4ex-KO model, where AQP4 staining surrounds the OMP^+^ OSNs (Fig. 2 a-c). We did not observe any change of AQP4 staining pattern in AQP4ex-KO (Fig. 2 a-c).

**Figure 2.**
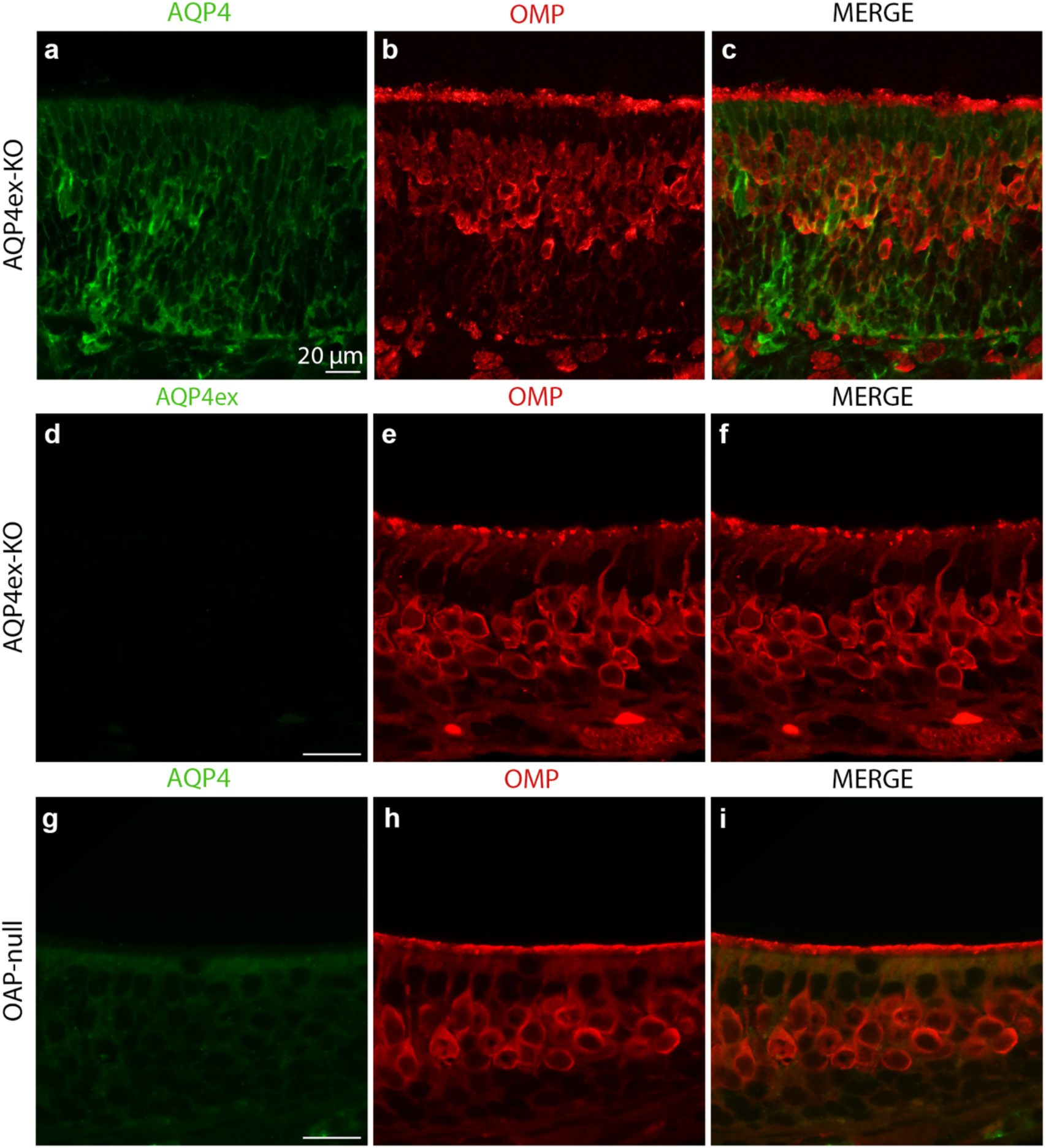
AQP4 localization is not affected by the lack of AQP4ex. (a, b, c) Confocal micrographs of coronal sections from AQP4ex-KO mouse olfactory epithelium showing AQP4 expression pattern. (d, e, f) AQP4ex staining is totally abolished in AQP4ex-KO, (g, h, i) and a sharp decrease of the staining is detectable in OAP-null referable to M1 isoform.

Finally, the weak and almost absent staining of AQP4 in OAP-null mice reveals that AQP4M23 isoforms- dependent OAP assembly is essential for AQP4 normal expression levels in the olfactory epithelium, as demonstrated in the brain (De Bellis et al., 2021) (Fig. 2 g-i).

### AQP4ex and AQP4M23 contribute to the structural integrity of the olfactory epithelium

While performing immunofluorescence experiments, we noticed that the cell density in the OE of AQP4ex- KO and OAP-null appeared sparser and reduced than in the WT. Therefore, we performed cell counting and found that mature OSNs, stained with OMP, were roughly 30% less in number in the KO models compared to WT (WT: 2524.55 ± 151.28 OMP^+^ cells/mm^2^ (mean cell density ± se), AQP4ex-KO: 1631.69 ± 59.14, OAP- null: 1529.76 ± 66.07) (Fig. 3 a, d). In addition, the SUSs, counted as SOX2^+^ nuclei in the apical part of the OE, were significantly lower in the KO models than WT (WT: 1008.93 ± 46.62 SUSs cells/mm^2^, AQP4ex- KO: 765.873 ± 56.10, OAP-null: 818.45 ± 50.84) (Fig. 3 b, d). The population of basal cells positive to SOX2 and Ki67, a maker for the GBCs, were also significantly lower in AQP4ex-KO and OAP-null compared to WT (WT: 213.29 ± 13.98 Ki67^+^ cells/mm^2^, AQP4ex-KO: 129.96 ± 16.29, OAP-null: 154.76 ± 20.96) (Fig. 3 c, e).

**Figure 3.**
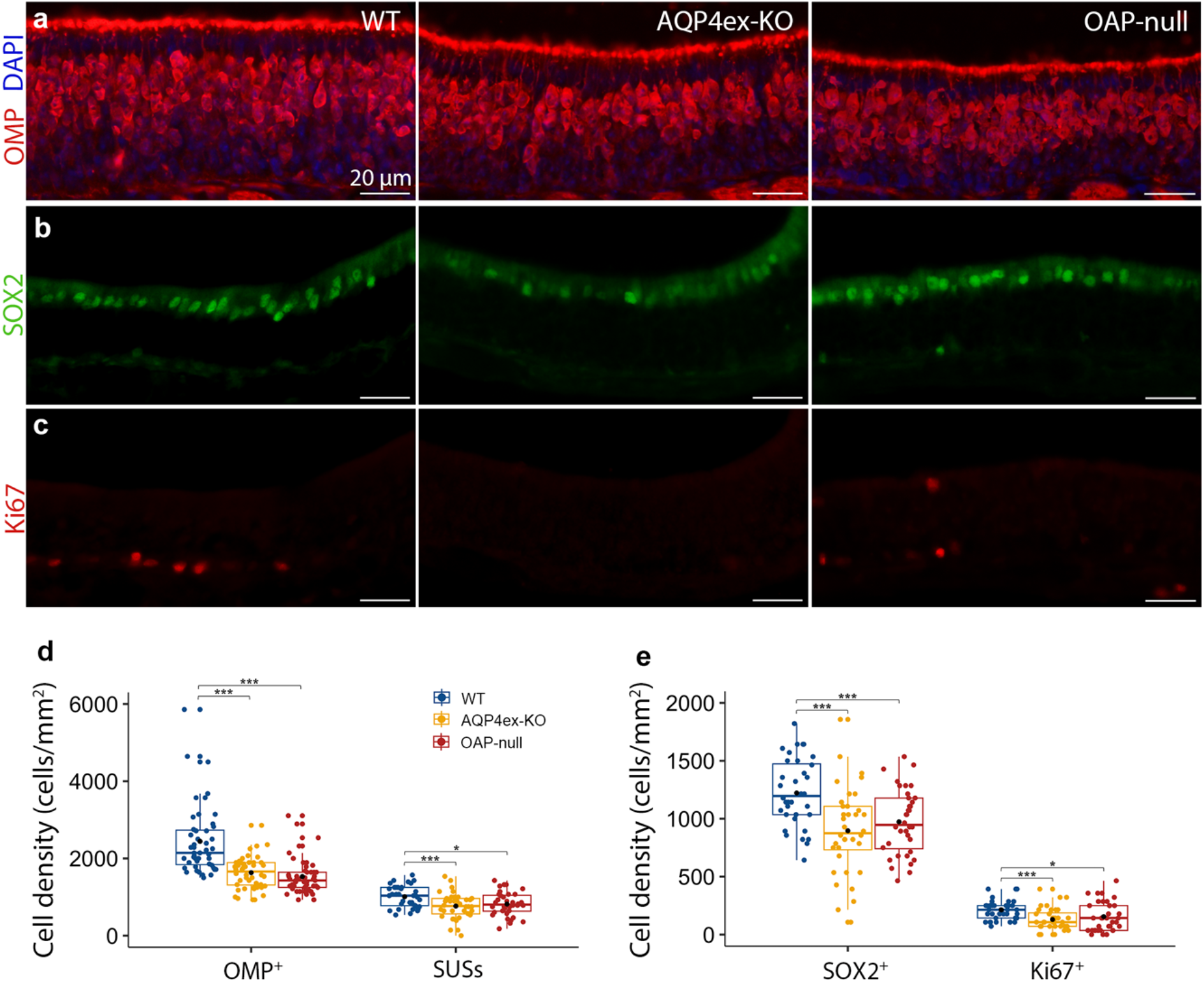
AQP4ex and AQP4M23 deletion affect cell density in the olfactory epithelium. (a) Coronal sections from olfactory epithelium showing OMP^+^ cells. (b, c) Coronal sections from olfactory epithelium showing SUS and GBC cells stained with Sox2 and Ki67 respectively. (a center, right; b center, right; d) Absence of AQP4ex and AQP4M23 reduces OMP^+^ and SUS cell density. (c center, right; e) At the same time, the cell density of GBCs is reduced in AQP4ex-KO and OAP-null. Box plots showing cell density in wild-type and knockout mice. The central point represents the mean, central line: median, upper and lower box boundaries: 25^th^ and 75^th^ percentile, extreme lines: the highest and lowest value (one-way Anova, followed by Tukey test post-hoc analysis, or Kruskal-Wallis test followed by Benjamini-Hochberg (BH) post-hoc analysis was performed. Anova I performed on SOX2^+^ (genotype: *F(*2, 105) = 10.34, *p* = 7.95e^-5^). Anova I performed on SUSs (genotype: *F(*2, 105) = 6.21, *p* = 3e^-3^). Kruskal-Wallis performed on Ki67^+^: *H*(2) = 13.39, *p* = 1.24e^-3^. Kruskal-Wallis performed on OMP^+^: *H*(2) = 52.71, *p* = 3.58e^-12^; *p* = * 0.05, ** 0.01, *** 0.001, *n* = 3 per genotype).

These findings were further supported by Western blot analysis. Here, we found that the percentage of OMP protein over the total protein quantity is decreased in the KO models compared to WT (Fig. 4 a center, b). Moreover, we could differentiate the bands corresponding to different AQP4 isoforms that highlight how AQP4ex is less abundant in the OE compared to the overall AQP4 and that AQP4M23 isoforms are more abundant than AQP4M1 (Fig. 4 a top). Densitometric analysis reveals that AQP4ex represents roughly 6% of the total AQP4 in the olfactory epithelium of WT tissue (Fig. 4 a bottom). Moreover, the absence of the AQP4ex does not substantially alter the percentage of other AQP4 isoforms (Fig. 4 b). In both KO mouse models, we did not find altered expression of Kir 4.1 (Fig. 4 a bottom, b), an inward rectifying potassium channel that seems mostly expressed in the microvilli of the SUSs (Fig. 4 c-e). Since in WB analysis we used the whole OE tissue, Kir 4.1 may be expressed in other cell types whose density does not change.

**Figure 4.**
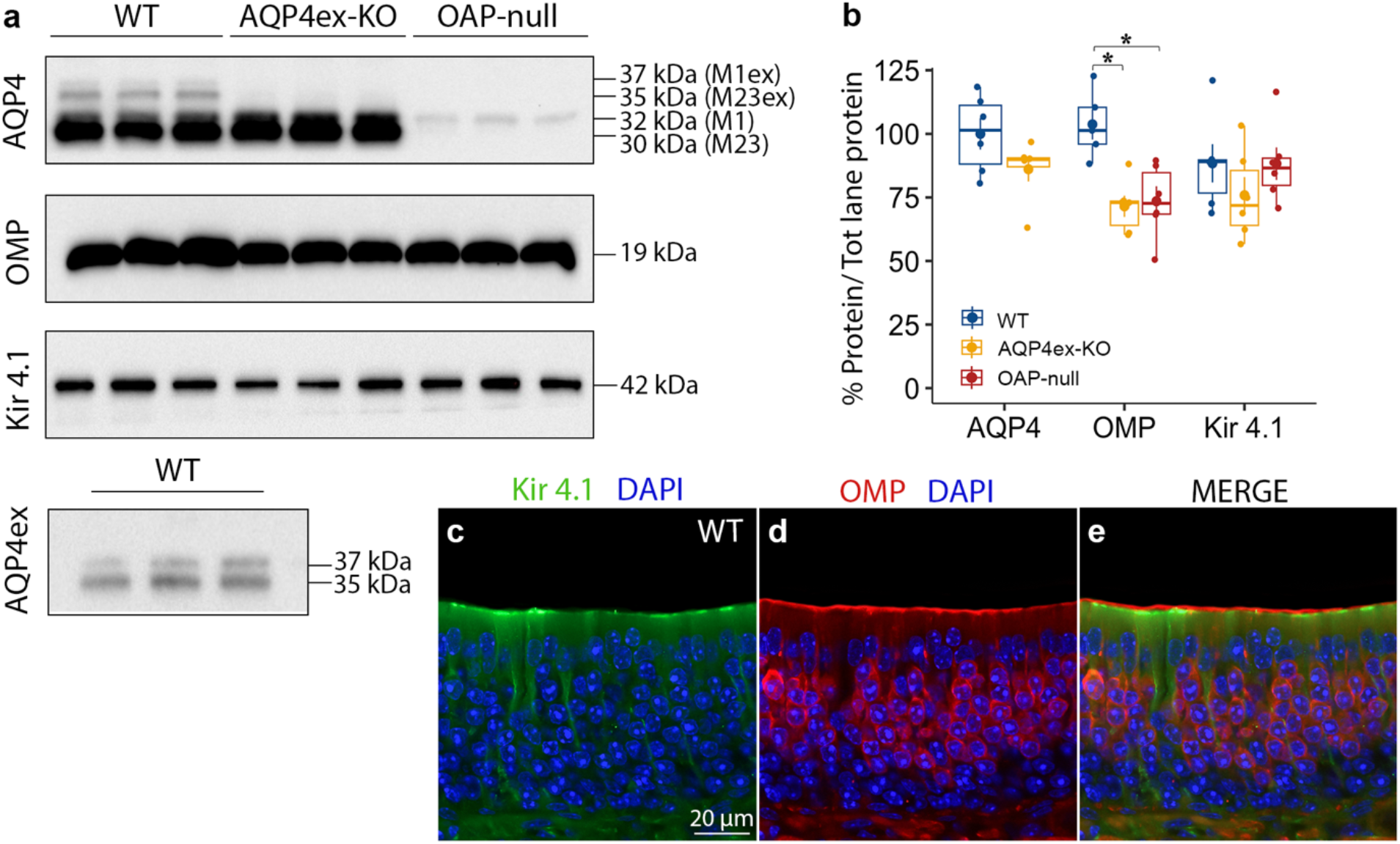
OMP expression is reduced in AQP4ex-KO and OAP-null. (a) Western blot analysis of olfactory epithelium proteins: AQP4, AQP4ex, OMP and Kir 4.1. Specimens probed with anti-AQP4 antibody, showing proteins at 30, 32, 35, 37 kDa corresponding to the four AQP4’s isoforms. (b) Box plots showing %AQP4, OMP and Kir 4.1 in wild-type and knockout mice. The central point represents the mean, central line: median, upper and lower box boundaries: 25^th^ and 75^th^ percentile, extreme lines: the highest and lowest value. Two-way Anova followed by Tukey test post-hoc analysis was conducted. Anova II performed on %protein in the OE revealed genotype as main effect (genotype: *F(*2, 39) = 7.32, *p* = 2e^-3^; *p* = * 0.05, ** 0.01, *** 0.001, *n* = 6 per genotype). (c, d, e) Confocal micrographs of coronal sections from wild-type mouse olfactory epithelium showing Kir 4.1 expression pattern localized on the microvilli of the sustentacular cells, below the cilia of the OSNs (OMP^+^ cells).

In summary, our results indicated that AQP4ex expression as well as the OAP-forming isoform AQP4M23 are necessary to maintain the proper ratio between the different cell types in the OE, suggesting that they are involved in setting a permissive micro-environment for cell development and differentiation.

### EOG responses are reduced in AQP4ex-KO and OAP-null mouse model

A direct consequence of the decrease in OSN density in the OE might be a reduction in the magnitude of the odorant response. A previous report described a decreased EOG response due to an altered coupling between water exchange and potassium clearance in the extracellular space of the OE (Lu et al., 2008). We performed air phase EOG recordings from the OE of WT, AQP4ex-KO and OAP-null mice. When stimulated with isoamyl acetate (IAA) at 10^−1^ M, we observed a response that was, on average, around -14 mV in WT and that decreased to about half in AQP4ex-KO and OAP-null mice (Fig. 5 a, b). The reduced response amplitude was neither odorant-dependent nor concentration-dependent. We stimulated the OE using a different odorant, geraniol (Fig. 5 b) and we could again observe a decrease in response amplitude in KO animals. Dose-response showed that the amplitude of the response at every tested concentration was smaller in the KO mouse models compared to the WT (Fig. 6 a, b), although we did not observe a clear shift in odorant sensitivity in both the KO. These results highlight the role of the AQP4 isoforms in maintaining OE functionality by ensuring and keeping the correct microenvironment for the OSNs. The lack of differences in EOG response kinetics further strengthens the idea that single cell OSN functionality could be unaltered in the AQP4ex-KO and OAP-null mice. Indeed, time to peak and decay time (*t*_*20*_) are similar across the three genotypes (Fig. 5 c, d).

**Figure 5.**
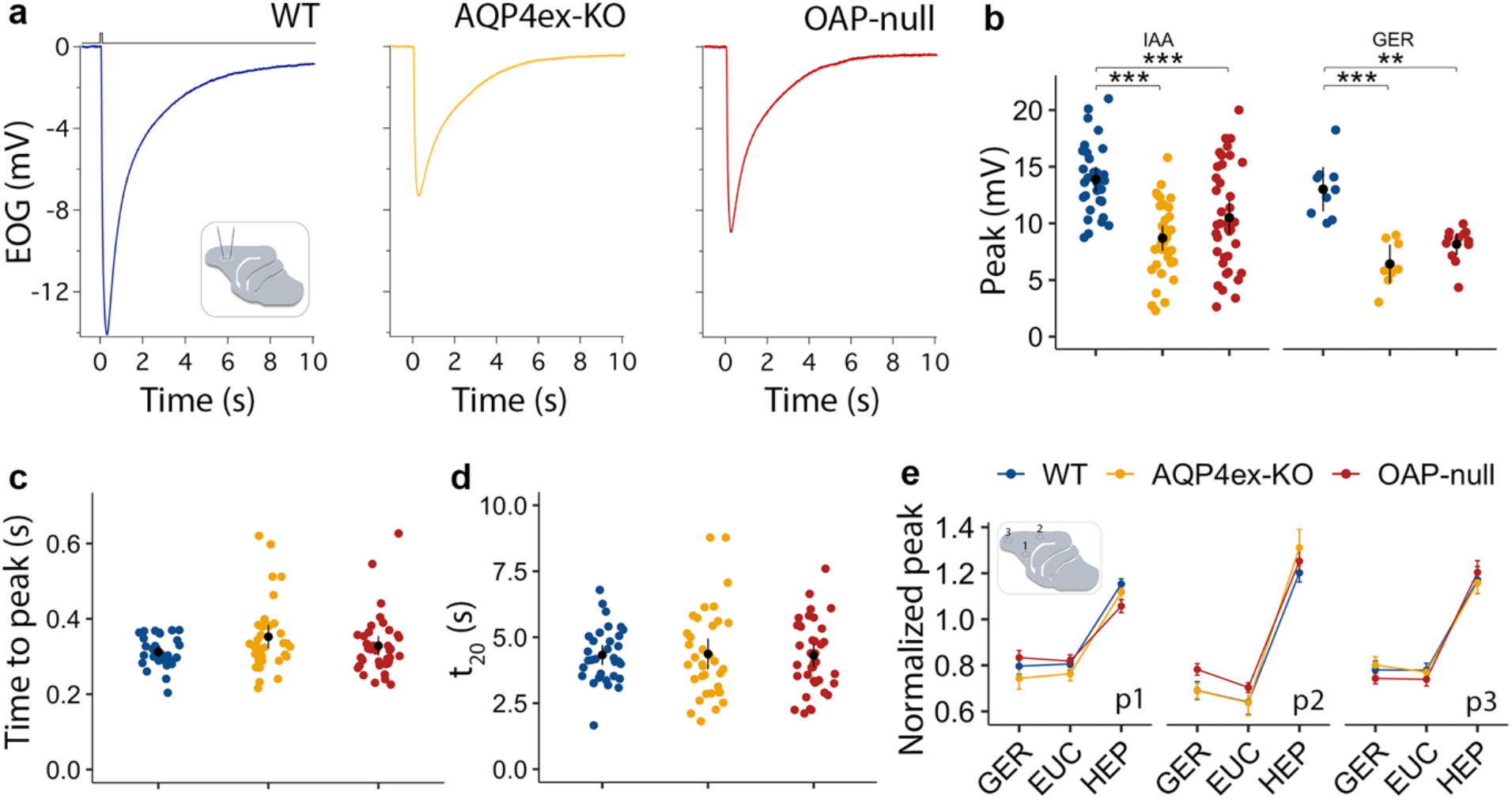
AQP4’s isoforms affect the odorant-evoked response. (a) Odorant-evoked EOG responses to 100 ms exposure to isoamyl acetate (10^−1^ M) were recorded from turbinate IIa of wild-type (blue), AQP4ex-KO (yellow) and OAP-null (red). (b) EOG amplitude response to 10^−1^ M isoamyl acetate and geraniol. EOG responses are reduced in AQP4ex-KO and OAP-null mice, data are presented as mean and ci. Two-way Anova followed by Tukey test post-hoc analysis was conducted. Anova II performed on EOG responses revealed genotype and odor as main effects (genotype: *F(*2, 128) = 27.18, *p* = 1.46e^-10^, odor: *F(*1, 128) = 6.58, *p* = 1.10e^-2^; *p* = * 0.05, ** 0.01, *** 0.001). (c, d) AQP4’s isoforms do not affect the kinetics of the response (data are presented as mean and ci. *t*_*20*_: 20% decay time, one-way Anova and Tukey test post-hoc analysis, *WT* = 41 mice in total, *AQP4ex-KO*= 43, *OAP-null* = 50). (e) EOG responses to 100 ms exposure to geraniol, eucaliptol, and heptaldehyde, all at 10^−1^ M were recorded from three positions (p1, p2, p3) of turbinate IIa. Peaks are normalized to isoamyl acetate, data are presented as mean ± se. Linear mixed model performed on normalized EOG responses revealed odorant as main significant effect (odorant: *F(*2, 190.3) = 353.19, *p* = 2.2e^-16^) and there were no differences between the three genotypes (*WT* = 6-9 mice, *AQP4ex-KO*= 6-9, *OAP-null* = 11-12). IAA: isoamyl acetate, GER: geraniol, EUC: eucaliptol, HEP: heptaldehyde.

**Figure 6.**
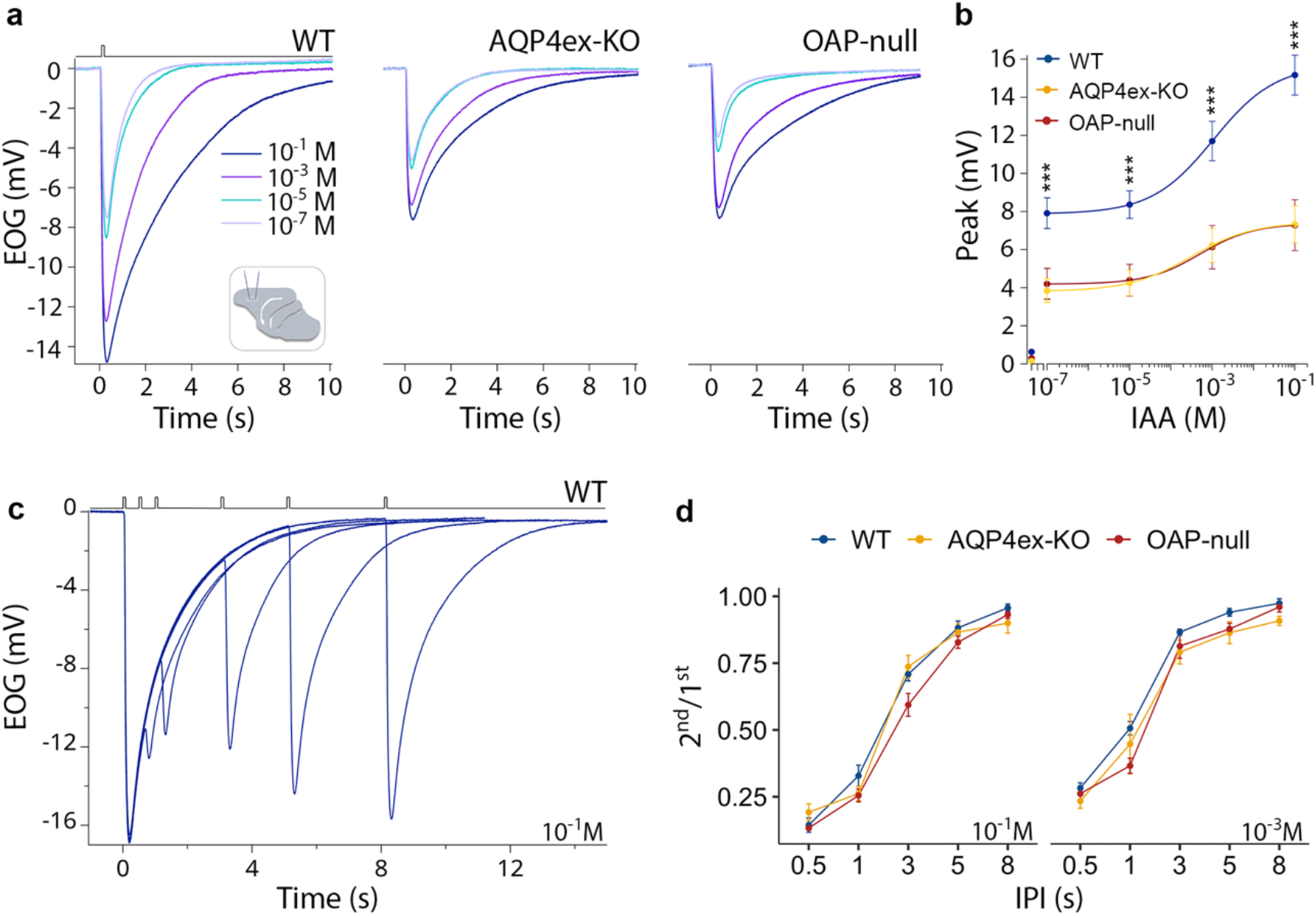
EOG responses from AQP4-KO mice, recover from adaptation similar to wild-type (a) Odorant-evoked EOG responses were evoked by 100 ms stimulation with isoamyl acetate vapor of increasing concentrations ranging from 10^−1^ to 10^−7^ M recorded from turbinate IIa of wild-type, AQP4ex-KO and OAP-null mice. (b) EOG amplitude responses are reduced in KO models. EOG responses to air are also shown; data are presented as mean ± se. Linear mixed model performed on EOG responses to different concentrations revealed genotype, dose and genotype-dose interaction as significant (genotype: *F(*2, 28.1) = 15.29, *p* = 3.22e^-5^, dose: *F*(3, 77.7) = 92.62, *p* = 2.2e^-16^, genotype-dose: *F*(6, 77.7) = 4.42, *p* = 6.80e^-4^; *p* = * 0.05, ** 0.01, *** 0.001), nonetheless, there is no difference in genotype-dose when EOG responses are normalized to the higher dose used. *WT* = 9 mice, *AQP4ex-KO*= 11, *OAP-null* = 9. (c) Paired pulse odorant responses evoked by 100 ms stimulation with different interpulse intervals (IPI: 0.5, 1, 3, 5, 8 s) to 10^−1^ M isoamyl a cetate (IAA) from wild-type. (d) Ratio of response to the second stimulus to the first ± sem at the indicated odorant concentration plotted versus the IPI; data are presented as mean ± se. Linear mixed model performed to 2^nd^/1^st^ response revealed ipi, dose and ipi-dose interaction as main significant effects (ipi: *F(*4, 149.9) = 787.94, *p* = 2.2e^-16^, dose: *F*(1, 153.3) = 69.49, *p* = 4.09e^-14^, ipi-dose: *F*(4, 149.9) = 7.54, *p* = 1.46e^-5^), *WT* = 7 mice, *AQP4ex-KO*= 6, *OAP-null* = 8.

In this scenario, the action of the AQP4 isoforms is maintaining the proper OE structural integrity rather than directly the OSN functionality.

A further confirmation that OSN response is not altered, comes from the paired-pulse paradigm experiments to explore adaptation. We applied a pulse of isoamyl acetate (IAA) of 100 ms each, with the time between pulses (interpulse intervals, IPI) varying from 0.5 to 8 s (Fig. 6 c, d). Since the response to the first odor pulse did not decay to baseline when the second pulse was applied, the recorded second response represents the sum of the residual first response and the adapted response to the second pulse. We found no significant differences across the genotypes.

Finally, we selected three odorants with different air-mucus odorant partition coefficients (see methods and Kurtz et al., 2004; Scott et al., 2014; Coppola et al., 2017), and found that we could still observe a significant reduction in the responses of the KO models compared to WT. Normalizing the responses to isoamyl acetate (IAA), hepthaldehyde (HEP) consistently elicited higher responses than eucalyptol (EUC) and geraniol (GER) in all models. Importantly, the response profile in KO mice was similar to WT (Fig. 5 e), suggesting that mucus water content likely remains unchanged in the KO models.

Altogether, these results show that the electrophysiological properties of OSNs remain intact, while the overall response amplitude is compromised due to the loss of functional support provided by the expression of AQP4 isoforms in SUS cells.

### Olfactory-driven behaviors are impaired in AQP4ex-KO and OAP-null mouse model

We sought to investigate how the observed changes combine to alter naïve behavior such as tracking odors to find hidden food. We performed the odor-guided food-seeking test consisting of a hidden piece of cookie under the cage bedding. The mouse is free to explore the cage, and the latency time to find the cookie is measured. This behavior is mainly dependent on the olfactory ability of the mouse. AQP4ex-KO and OAP-null took more time to find the cookie than WT. Three over fifteen AQP4ex-KO and OAP-null mice did not find the cookie within 5 min and were excluded. Instead, all the tested WT mice found the cookie (Fig. 7 a). Interestingly, while we did not find differences in molecular and electrophysiological properties between AQP4ex-KO and OAP-null model, in the cookie test, we found that the former was slower than the latter. We did not find differences across the genotypes when the cookie was placed on top of the bedding, hence the animal could easily reach it and not have to rely on olfaction to find the cookie.

**Figure 7.**
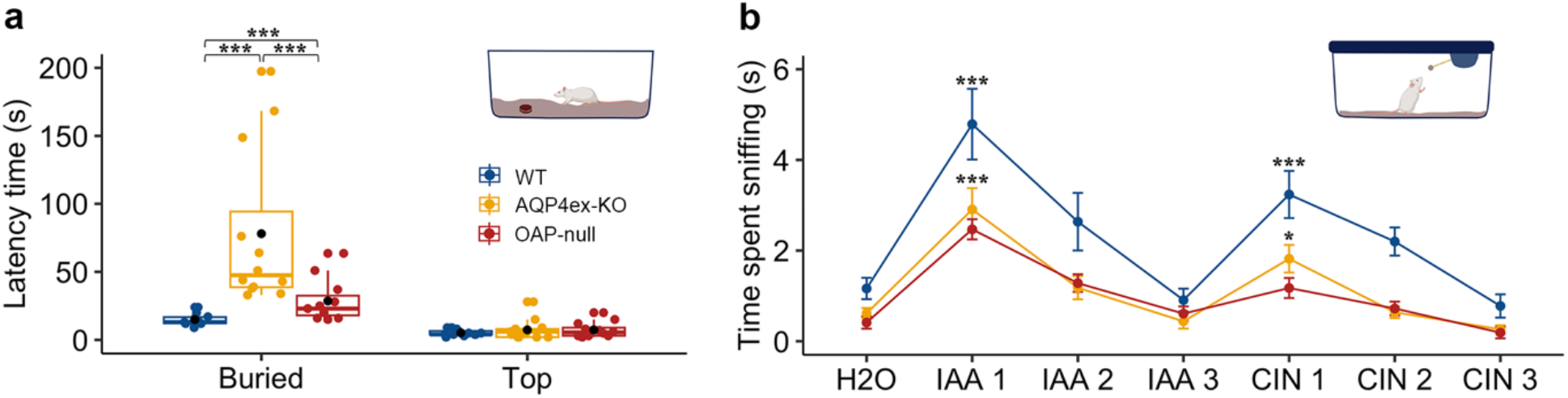
AQP4 isoforms affect olfactory abilities and discrimination. (a) Latency time in finding the cookie buried, or placed on top of the bedding. AQP4ex-KO and OAP-null mice are slower than wild-type in finding the cookie. The central point in the box plots represents the mean, central line: median, upper and lower box boundaries: 25^th^ and 75^th^ percentile, extreme lines: the highest and lowest value. Kruskal-Wallis test followed by Benjamini-Hochberg (BH) post-hoc analysis was performed on buried cookie: *H*(2) = 21.74, *p* = 1.91e^-5^; *p* = * 0.05, ** 0.01, *** 0.001, *WT* = 10 mice, *AQP4ex-KO* = 12, *OAP-null* = 12. (b) Time spent sniffing the new odorant within 2 min after the odorant previously presented. IAA and CIN were used at 1:100. Linear mixed model revealed genotype and odorant as main significant effects (genotype: *F(*2, 72.1) = 30.71, *p* = 4.354 e^-14^, odorant: *F*(6,182.8) = 30.77, *p* = 2.2 e^-16^). The statistical differences are to be interpreted as the difference between H2O and IAA1, and IAA3 and CIN1 for each genotype. There are no differences between IAA3 and CIN1 in OAP-null, instead AQP4ex-KO is different from wild-type (*p* = * 0.05). Data are presented as mean ± se, *WT* = 10 mice, *AQP4ex-KO* = 10, *OAP-null* = 10. IAA: isoamyl acetate, CIN: 1,4 cineole.

We also analyzed mouse behavior by using the habituation/dishabituation test. Habituation is the process whereby the mouse response to a stimulus decreases with repeated exposures, representing non-associative learning. We presented IAA to the mouse, and this step was repeated three times, and for each iteration, the length of time that the animal spent sniffing the odor source was noted. A reduction in sniffing time over successive trials is interpreted as evidence of the animal recognizing the odor. To assess odor dishabituation, we introduced cineole (CIN) at the fourth trial. The observed increase in investigation time indicates the ability of the mouse to differentiate between the previous (IAA) and new odor (CIN). The three mouse models we tested, adapted to the repeated exposures to IAA and CIN. Although the AQP4ex-KO mice showed a reduced time spent sniffing the new odor (CIN 1) compared to the WT, they increased their investigation time when the new odor was presented. Instead, the OAP-null mice failed to recognize the new odor, as indicated by the lack of difference in investigation time between the odor they had already sniffed (IAA 3) and the new odor (CIN 1) (Fig. 7 b). In summary, we could dissect how structural and functional impairments affected olfactory behavior by finding differences between AQP4ex-KO and OAP-null. The former was the slowest in finding the cookie, and the latter was unable to discriminate between different odors.

## Discussion

The role of SUS cells in the OE has long been neglected even though they are known to perform various functions, ranging from absorption, detoxification, metabolism, nourishment, phagocytosis, physical support and secretion (Chen et al., 1992; Getchell and Getchell, 1992; Suzuki et al., 1996; Hansen et al., 1998; Hegg et al., 2003). SUS cells in the OE abundantly express AQP4 (Fig. 1, 2; Lu et al., 2008). Similarly, AQP4 is widely expressed in sensory systems, such as the inner ear, retina, and vomeronasal organ (Ablimit et al., 2008; Sakai et al., 2014; Maurya et al., 2015; Amann et al., 2016). AQP4 is expressed in the organ of Corti, where the protein seems to play an essential role in inner ear fluid dynamics (Eckhard et al., 2012; Morris et al., 2012; Gleiser, et al., 2016). Here, we investigated for the first time the expression of the different AQP4 isoforms and their contribution to the OE physiology.

### AQP4ex and AQP4M23 isoforms ensure the structural and functional integrity of the olfactory epithelium

Since AQP4 can undergo different translational modifications, various isoforms of the protein exist, and based on the selected translational start point, AQP4M1 or AQP4M23 can be produced (Neely et al., 1999; De Bellis et al., 2017). The two isoforms exhibit a similar water permeability, even though the aggregation to generate OAPs and cellular distribution in the glial cells are different (Crane et al., 2008; Ciappelloni et al., 2019). For instance, AQP4M23 and AQP4M1 organize in OAP of varying sizes, resulting in an increased water permeability (Silberstein et al., 2004; Crane et al., 2009; Rossi et al., 2010). Recent studies shed light on the link between K^+^ uptake by astrocytes and water permeability through AQP4. A hypothesis of the K^+^-water coupling is that K^+^ uptake by astrocytes following neuroexcitation results in osmotic water influx, then producing astrocytes swelling and ECS shrinkage, increasing the K^+^ concentration and accelerating its uptake (Amiry-Moghaddam et al., 2003; Binder et al., 2006; Strohschein et al., 2011; Jin et al., 2012). Hence, astrocytes can maintain an optimum ionic microenvironment in the ECS through AQP4 that organizes in OAP, by controlling the water balance and allowing the neurons to maintain the right ion concentration balance for neuroexcitation. AQP4M23 is particularly crucial in guaranteeing all these mechanisms since, as previously described, the kinetics of the water permeability is modified when tested in primary cultures of astrocytes where AQP4M23 is missing (Solenov et al., 2004; De Bellis et al., 2021).

As already demonstrated in the brain (De Bellis et al., 2021), here we show that AQP4 staining is almost absent when AQP4M23 is missing, suggesting that the aggregation of AQP4 into OAPs is crucial for maintaining normal AQP4 expression levels and AQP4M1 alone is not sufficient to maintain the AQP4 channel localization on the basolateral membrane of the SUSs. Therefore, these cells cannot rely on AQP4 tetramers as an alternative to OAPs. The proper supramolecular organization would be important either to maintain the correct extracellular volume microenvironment or to allow the cell to structurally adapt to the ever-changing conditions of the OE.

AQP4ex isoform is expressed at the perivascular end-feet of the astrocytes (Palazzo et al., 2020) and it is necessary for perivascular localization of AQP4 (Palazzo et al., 2019; Palazzo et al., 2020) and BBB integrity (Mueller et al., 2023), along with α-syntrophin that is necessary to anchor AQP4 to the perivascular basal lamina (Neely et al., 2001; Amiry-Moghaddam et al., 2003). Therefore, AQP4 is no longer localized in the astrocyte end-feet facing the blood vessels in AQP4ex-KO (Palazzo et al., 2019). In the OE the lack of the extended isoform does not seem to affect the expression and localization of AQP4, suggesting that AQP4ex has a different role in the OE than in the brain (Palazzo et al., 2019). Although it has been shown that the lack of AQP4 did not impair OE morphology (Lu et al., 2008), detailed cell counting, as well as isoform-specific characterization, was not directly attempted. Since EOG measures the summated generator potential from the responding OSNs, the reduced OSN density observed in both transgenic mice would explain the reduced amplitude of EOG responses, without affecting the kinetics of the odorant response. These results suggest the involvement of AQP4ex and OAPs in preserving OE integrity rather than a direct contribution to the OSN transduction machinery. Furthermore, AQP4 most likely does not contribute to setting the composition of the mucus by altering its water content in the OE since odorants with different mucus solubility have similar responses in AQP4ex-KO and OAP-null. The basolateral localization of AQP4 in the SUSs further supports this idea.

The immunostaining reveals that AQP4ex isoform is most likely expressed in the OECs beneath the basal lamina in the OE. This cell type seems to be involved in roles such as olfactory development, phagocytosis, and in responding to neuroinflammation by reducing the microglia activation and pro-inflammatory factors release, thereby ameliorating the detrimental condition of the microenvironment (Lipson et al., 2003; Pastrana et al., 2006; Leung et al., 2008; Rojas-Mayorquín et al., 2008; Doncel-Pérez et al., 2009; Zhang et al., 2021). We reasoned that a role the OECs are involved in, might be mediated by AQP4ex that favors AQP4 localization in those cells, which may provide structural support and extend OECs processes that wrap the axons, while SUS cells do it for OSN cells body and dendrite, aiding in the formation of connections with the OB. Therefore, OECs lacking AQP4ex could fail to properly guide OSNs to their final target, thus causing OSN apoptosis, which would also explain the reduction in the OSNs number we observe.

### AQP4ex and AQP4M23 isoforms impact olfactory detection and discrimination

The odor-guided food-seeking test is a naïve behavioral test used to test the olfactory ability of mice to find hidden sources of food under the cage bedding. It is widely used, and the time the mouse takes to locate the food is dependent on a functionally intact olfactory system. The contribution of the peripheral olfactory system in such behavior is crucial since altered transduction in the OSNs alters the test (Klein et al., 1996; Pietra et al., 2016). Since the test does not rely on an operant-conditioning task, mice must use their innate sense of smell to locate the odor source food. AQP4 knock-out mice have been shown to find the food later than WT (Lu et al., 2008), and here we extended this finding by showing that both AQP4ex-KO and OAP-null mice are slower than WT, suggesting that both the isoforms are crucial to maintaining olfactory abilities, as expected from the EOG results. We further dissected the altered behavior since we found differences between the two transgenic mouse models. AQP4ex-KO showed a reduced ability to detect the odor source than OAP-null. Nonetheless, the amplitude of EOG response is similar in both knock-out mouse models, suggesting that olfactory processing could be altered in the olfactory bulb or higher olfactory circuitries. Hence, a different behavior may indicate that further along in olfactory processing in the brain, other changes occur that may globally explain the different behavior we observe.

The habituation test proved that both AQP4ex-KO and OAP-null mouse models could adapt similarly to WT. Interestingly, after behavioral habituation, OAP-null failed to dishabituate, and then spent less time than WT and AQP4ex-KO mice to investigate the new odorant.

A possible explanation of this behavior is that the altered water dynamics could modify the extracellular space and probably basolateral ionic homeostasis. While this alteration may not significantly affect ciliary olfactory transduction, it could impair the firing of action potentials.

The use of the two different animal models gave interesting insights about the expression and localization of AQP4 in the OE. In the OAP-null mouse model, the absence of OAPs results in a marked reduction of AQP4, effectively creating a similar AQP4 knockout condition. Despite AQP4ex-KO maintaining physiological levels of AQP4 in the OE, without exhibiting altered membrane organization, both models exhibit similar detrimental effects on the OE, such as decreased cell density and impaired olfactory functions.

Overall, we showed that the contribution of the SUSs in preserving the microenvironment surrounding OSNs and the other cell types of the OE could be modified when one of the AQP4’s isoforms is missing. Finally, we sought to investigate how the two isoforms impact behavior, and their contribution to olfactory detection and discrimination, finding that olfactory abilities are impaired. Hence, our results establish a foundation to further understand the precise involvement of the two isoforms in the OB and other olfactory circuitries.

## Methods

### Animals

C57BL/6J OAP-null and AQP4ex KO mice were generated by Cyagen Biosciences Inc. (Santa Clara, USA) using CRISPR/Cas9 technology as previously described (Palazzo et al., 2019; De Bellis et al., 2021). Mice were maintained under a light/dark cycle (12/12 h) in the Department of Translational Biomedicine and Neuroscience’s animal facility at room temperature (22±2°C) and 75% humidity, with food and water ad libitum. All experiments were performed using procedures approved by the Institutional Committee on Animal Research and Ethics of the University of Bari and the Italian Health Department (Project n°475/2020-PR) and in accordance with the European directive on animal use for research. Experiments were carried out on mice from 1 to 6 months old and of either sex. All experiments were designed to minimize the number of animals used and their suffering.

### Immunohistochemistry

Mice (1 month old) were anaesthetized with CO_2_ inhalation and decapitated. The dissected nose was fixed in 4% PFA solution overnight at 4°C and washed in PBS Ca^2+^-Mg^2+^, pH 7.4. Noses were decalcified in EDTA (0.5 M, pH 8) over several days up to a week at 4°C with sporadic replacement of the buffer. The time of decalcification depends on the degree of mineralization of the bone. After washing in PBS Ca^2+^-Mg^2+^, the buffer solution was replaced with sucrose at 5, 10, and 20% (w/v) in PBS Ca^2+^-Mg^2+^ for 10 min each. Then, tissues were placed in 30% (w/v) sucrose, overnight at 4°C. After washing in the buffer solution, tissues were frozen in optimal cutting medium compound Tissue-Tek OCT (Sakura, The Netherlands). Coronal sections 14-15 μm thick were cut with a cryostat (CM 1900, Leica) at -20°C and stored on poly-L-lysine glass slides at -80°C. After rehydration with PBS Ca^2+^-Mg^2+^, sections were incubated with SDS 0.5% (w/v) in PBS Ca^2+^- Mg^2+^ for antigen retrieval, followed by blocking solution 5% (v/v) normal goat serum, 4% (w/v) BSA, 5% (w/v) non-fat dry milk, 0.3% (w/v) Triton X-100 in PBS Ca^2+^-Mg^2+^, for 30 min. Then the slices were incubated overnight at 4°C with rabbit polyclonal anti-AQP4 (GenScript Biotech, Piscataway, NJ, USA) diluted 1:2000 in the blocking buffer, rabbit polyclonal anti-AQP4ex generated against the peptide DSTEGRRDSLDLASC within the AQP4 C-terminus (1:1000; GenScript Biotech, Piscataway, NJ, USA), mouse monoclonal anti-OMP (1:300; Santa Cruz Biotechnology, TX, USA), rat monoclonal anti-Sox2 (1:300; eBioscience, Invitrogen, San Diego, USA), rabbit polyclonal anti-Kir 4.1 (1:200, Alomone Labs, Jerusalem, Israel), rabbit polyclonal anti-Ki67 (1:200; Abcam, Cambridge, UK). AlexaFluor 488 goat anti-rabbit, AlexaFluor 594 donkey anti-rabbit, and AlexaFluor 594 goat anti-mouse were used at 1:1000, and AlexaFluor 488 donkey anti-rat (1:500) in 0.2% (w/v) Tween 20, PBS Ca^2+^-Mg^2+^ (all from Life Technologies, Thermo Fisher Scientific, Carlsbad, California, USA) for 45 min at room temperature. After washing with PBS Ca^2+^-Mg^2+^, sections were mounted with 1:1 Mowiol (Sigma-Aldrich, Waltham, MA, USA) - DAPI (Life Technologies, Thermo Fisher Scientific, Waltham, MA, USA). Images were acquired with a confocal laser scanning microscope (TCS SP8, Leica) using 40x/1.30 or 63x/1.40 HC PL Apo CS2 oil objective at 1024 × 1024p and analyzed with Fiji software (Schindelin et al., 2012).

### Cell counting

The number of cells with immunoreactive signal for OMP, Sox2, and Ki67 were counted from 12 areas of coronal sections 15 μm thick, spanning widely separated regions in the olfactory epithelium from the anterior part of turbinate II to the ventral region of turbinate IV. Sustentacular cells were counted, excluding Sox2^+^ stem cells laying on the basal lamina (*n* = 3 mice, 5-6 months old). Cell density was measured in a 117 (h) x 236 (w) μm^2^ area, that included all the OE thickness. Images were acquired with an LED fluorescence microscope (DM2500, Leica) using 20X/0.55 HC PL FLUOTAR objective at 1280 × 1024p and a CCD camera (DFC7000-T, LEICA), then analyzed with Fiji software.

### SDS-PAGE and Western blotting

Mice (3-6 months old of either sex) were anaesthetized with CO_2_ inhalation and decapitated. The head was split midsagitally, and the olfactory epithelium and bulb were removed. The specimens were stored into liquid nitrogen. Samples were dissolved in BN buffer (5-7x of the sample volume) (1% Triton X-100 (w/v), 12 mM NaCl (w/v), 500 mM 6-aminohexanoic acid (w/v), 20 mM Bis-Tris (v/v) pH 7.0, 2 mM EDTA (w/v), 10% glycerol (w/v)) and a Protease Inhibitor Cocktail (Roche, Milan, Italy). Lysis was performed on ice, vortexing the samples every 5 min for 30 min, and then the specimens were centrifugated at 15000 rpm at 4°C for 45 min. The supernatant was collected and stored at -80°C, and the protein concentration was measured with a BCA Protein Assay Kit (Thermo Scientific, Waltham, Massachusetts). The electrophoresis and immunoblotting were performed as previously described (Nicchia et al., 2009). Briefly, proteins were separated on 13% acrylamide/bis-acrylamide gel for AQP4 (20 μg), AQP4ex (20 μg) and Kir 4.1 (20 μg) and 15% acrylamide/bis-acrylamide gel for OMP (35 μg), and transferred to polyvinylidenedifluoride (PVDF) membranes (Millipore, Burlington, Massachusetts, USA) for immunoblot analysis. Membranes were incubated overnight at 4°C with the following primary antibodies: rabbit polyclonal anti-AQP4 (1:4000, GenScript Biotech, Piscataway, NJ, USA), rabbit polyclonal anti-AQP4ex (1:2000; GenScript Biotech, Piscataway, NJ, USA), mouse monoclonal anti-OMP (1:1000; Santa Cruz, Biotechnology, TX, USA), rabbit polyclonal anti-Kir 4.1 (1:400, Alomone Labs, Jerusalem, Israel). Then washed and incubated with the following peroxidase-conjugated secondary antibodies at room temperature for 45 min: goat anti-rabbit IgG HRP (1:3000; Bio-Rad, California, USA) and goat anti-mouse IgG HRP (1:3000; Bio-Rad, California, USA). Proteins were detected using an enhanced chemiluminescent detection system (Clarity Western ECL Substrate, Bio-Rad, California, USA) and visualized with Chemidoc Touch imaging system (Bio-Rad, California, USA). Densitometry analysis was performed using Image Lab (Bio-Rad, California, USA).

### Electro-olfactogram recording

The experimental procedure is similar to that previously described by Zhao et al., 1998, Cygnar et al., 2010 and Guarneri et al., 2023. Mice (5-6 months old) were anaesthetized with CO_2_ inhalation and decapitated. The head was split sagittally along the midline and immediately placed on a dissecting microscope (Stemi DV4, Zeiss) on a vibration-isolating table (Supertech Instruments) and shielded from electrical fields by a Faraday cage. The turbinates were exposed by removing the septum, and the specimen was continuously perfused with bubbled distilled water at 37°C to guarantee the hydration of the epithelium. The recording electrode was made from borosilicate glass (World Precision Instruments) pulled with a P-1000 puller (Sutter Instrument) and then placed on the surface of one of the turbinates. The pipette was filled with Ringer’s solution (mM): 140 NaCl, 5 KCl, 2 CaCl_2_, 1 MgCl_2_, 10 Hepes, 0.01 EDTA, pH 7.4 with NaOH, and the tip of the pipette was filled with 0.5% (w/v) agar in Ringer. The stimulus pulse was controlled by a Picospritzer-controlled valve (Picospritzer II, Parker Hannifin). A 100 ms pulse of vapor phase odorant was injected at 10 psi into a continuous stream of humidified air. A flow meter (Masterflex) set to 3 L/min was used to control the flow of humidified air arriving at the sample. The device connected the Picospritzer to the delivery tube. The odorant vapor was generated by placing an odorant solution in a 5 ml glass test tube sealed with a rubber stopper. The system used two 20- gauge needles as input and output ports of the vapor above the solution. The reference electrode was electrically connected to the tissue by positioning into the mouse brain. The odorants used: isoamyl acetate (IAA), geraniol (GER), eucalyptol and heptaldheyde, were prepared every day from 5 M stock in dimethyl sulfoxide (DMSO) (Honeywell, Riedel de Haën, Seelze, Germany), then diluted in water to reach a final concentration from 10^−7^ to 10^−1^ M. The log (air/mucus partition coefficients) were obtained from (Kurtz et al., 2004; Scott et al., 2014; Coppola et al., 2017) and were -2.4038 for isoamyl acetate, -2.766 for heptaldheyde, -3.4368 for eucalyptol and -5 for geraniol.

EOG responses were recorded at room temperature. Data were collected with an EPC 10 USB (HEKA, Elektronik) amplifier and analyzed using PatchMaster Next (version 1.4.1). The signals were recorded at a sampling rate of 10 kHz and low-pass filtered at 2.9 kHz. Kinetics of EOG responses were evaluated, measuring time to peak, decay time, rise time, latency time and FWHM (full width at half maximum) (data not shown). Time to peak was determined as the time from the start of the stimulation to the peak of the response; decay time (*t*_*20*_) as the time the response takes to decrease to 20% of the peak; latency as the time between the start of odorant stimulation to 1% of the peak value; the rise time as the time between the start of the response and the peak. All chemicals were purchased from Sigma-Aldrich.

### Odor-guided food-seeking test

A small piece of food was buried below the surface of the cage bedding so that it was a purely olfactory cue, and the time taken by the mouse to find the food was recorded. All mice of either sex (5-6 months old) were maintained under a light/dark cycle (12/12 h) and tested during the light phase. Mice were food-deprived overnight with ad libitum access to water. Before the test, each mouse was familiarized with the experimental setting by placement in the enclosure for 5 min, and then placed in a novel cage (l x h x w; 30 cm x 12 cm x 15 cm) where a ∼3 g piece of cookie (Oreo) had been buried under ∼2 cm of fresh bedding. The latency of the animal to retrieve the cookie was recorded with a stopwatch. Retrieving the cookie was defined as digging it up with forepaws, picking it up, and placing it in the mouth. We set a 5 min limit within which the animals had to find the cookie. After that, the time was stopped, and the cookie was exposed so the mouse could access and eat it. The position of the buried cookie was randomly changed in each experiment. The next day, the cookie was placed on the bedding. Since just some KO mice could not find the cookie within 5 min, we excluded them.

### Habituation/dishabituation test

A progressive decrease in olfactory investigation time towards a repeated presentation of the same odor stimulus defines habituation. Dishabituation allows the reinstatement of odor investigation when a novel odor is presented (Woodley and Baum, 2003; Wrenn et al., 2004). Therefore, this test assesses if the animal is able to distinguish different odors. The experimental procedure is similar to that previously described by Yang et al., 2009. Briefly, before the test, each mouse was acclimated for 30 min in a dedicated room and placed in a novel cage (l x h x w; 30 cm x 12 cm x 15 cm) where a clean and dry applicator was inserted through the water bottle hole of the cage lid. This procedure is crucial to reduce novelty-induced exploratory activity during the olfactory test. Then, each mouse was exposed to sequential presentations of different odors in the following order: water, isoamyl acetate (IAA) and geraniol (GER) diluted 1:100 in water. Each odor was presented in three consecutive trials for 2 min and the time spent sniffing the applicator with the odorant was recorded with a stopwatch. The inter-trial interval was 1 min. The cotton tip part of the applicator did not touch any part of the cage lid while changing the odor, avoiding cross-trial contamination.

### Statistical analysis

Statistical analysis was conducted using R (R Studio, 2024). Sample distribution was assessed for normality with Shapiro-Wilk test using *shapiro*.*test()* function, and parametric or non-parametric test conducted accordingly. Anova (*anova_test()*) from *“rstatix”* package (Kassambara, 2023), followed by Tukey test post-hoc analysis was used as the parametric test. Kruskal-Wallis (*kruskal*.*test()*), followed by Benjamini-Hochberg (BH) post-hoc analysis (*pairwise*.*wilcox*.*test()*) was used as a non-parametric test. All the aforementioned functions were used by running *“stats”* package (R Core Team, 2017). The *“emmeans”* package (Russell, 2023) was used for performing pairwise comparisons of estimated marginal means (EMMs) by using *pairs()* function. EMMs compare the levels of a factor by removing the effects of other factors, and were used to interpret the main effect and interactions. Mixed model Anova was used for multilevel analyses. In particular, in R environment, dependent variables were investigated with linear mixed models (LMMs). LMMs were computed using the *lmer()* function (*“lme4”* package, Bates et al., 2015). Finally, in order to get *F* statistics and *p*-value for the fixed effects of the models, we ran Anova using *“lmerTest”* package (Kuznetsova et al., 2017). Post-hoc comparisons were performed as stated in the figure legends. *“ggplot2”* package (Wickham, 2016) was used to create the graphs.

## Additional information

The data that support the findings of this study are available from the first and corresponding authors upon reasonable request.

## Competing interests

None declared.

## Author contributions

DL performed electrophysiological and behavioral experiments. DL and PA performed Western Blotting. DL and MGF performed immunofluorescence experiments. DL, MGF and BB performed image acquisition, DL and MD analyzed the experiments. PA, OV, CP collection of initial data.

MD, DL, AF and GPN conceptualized and wrote the manuscript. All the authors approved the final version of the manuscript.

## Fundings

The research was supported by AstroDyn (FA9550-19-1-0370) and AstroColl (FA9550-21-1-00352) funded by AFOSR to GPN; Stochastic Biophysical Interactions within Aquaporin-4 Assemblies (FA9550-20-1-0324) funded by AFOSR to GPN; Marie Skłodowska-Curie Actions -ITN-2020 ASTROTECH (GA956325) funded by the European Commission to GPN; project CN00000041 - National Center for Gene Therapy and Drugs based on RNA Technology (DD n.1035, 17.06.2022) and project MNESYS (PE0000006, DD n.1553, 11.10.2022) both from the National Recovery and Resilience Plan (PNRR), to GPN and AF. NANODYN PRIN*–Bando 2022 PNRR (P2022Z27NS) to GPN*. PRIN 2022 project 202297W2H3 - “Odor Coding and Transduction in human olfactory sensory neurons” to MD. Investment PE8 – project Age-It: “Ageing Well in an Ageing Society” (DM n. 1557, 11.10.2022) to AF and PRIN 2022 to AF.

## Acknowledgements

We thank Dr. Alessia Ricci for technical help with the building of the EOG set-up. We thank Dr. Johannes Reisert for very helpful advices and support.

## Notes

### Competing Interest Statement

The authors have declared no competing interest.

